# Insulin sensitization by hepatic FoxO deletion is insufficient to lower atherosclerosis in mice

**DOI:** 10.1101/2023.10.14.562366

**Authors:** María Concepción Izquierdo, Michael Harris, Niroshan Shanmugarajah, Kendra Zhong, Lale Ozcan, Gabrielle Fredman, Rebecca A. Haeusler

**Affiliations:** Naomi Berrie Diabetes Center, Columbia University College of Physicians and Surgeons; New York, NY, 10032; USA; Department of Pathology and Cell Biology, and Columbia University College of Physicians and Surgeons; New York, NY, 10032; USA; Department of Medicine; Columbia University College of Physicians and Surgeons; New York, NY, 10032; USA; Department of Molecular and Cellular Physiology, Albany Medical College, Albany, NY, 12208; USA

## Abstract

**Background:** Type 2 diabetes is associated with an increased risk of atherosclerotic cardiovascular disease. It has been suggested that insulin resistance underlies this link, possibly by altering the functions of cells in the artery wall. We aimed to test whether improving systemic insulin sensitivity reduces atherosclerosis.

**Methods:** We used mice that are established to have improved systemic insulin sensitivity: those lacking FoxO transcription factors in hepatocytes. Three hepatic FoxO isoforms (FoxO1, FoxO3, and FoxO4) function together to promote hepatic glucose output, and ablating them lowers glucose production, lowers circulating glucose and insulin, and improves systemic insulin sensitivity. We made these mice susceptible to atherosclerosis in two different ways, by injecting them with gain-of-function AAV8.mPcsk9^D377Y^ and by crossing with Ldlr^−/−^ mice.

**Results:** We verified that hepatic FoxO ablation improves systemic insulin sensitivity in these atherosclerotic settings. We observed that FoxO deficiency caused no reductions in atherosclerosis, and in some cases increased atherosclerosis. These phenotypes coincided with large increases in circulating triglycerides in FoxO-ablated mice.

**Conclusions:** These findings suggest that systemic insulin sensitization is insufficient to reduce atherosclerosis.

**Non-standard Abbreviations and Acronyms:** Ldlr, low-density lipoprotein receptor; TRL, triglyceride-rich lipoprotein; VLDL, very low-density lipoprotein

## Introduction

Individuals with type 2 diabetes and its forerunner, the metabolic syndrome, are at an increased risk of atherosclerotic cardiovascular disease^1^. A potential explanation for this tight link is insulin resistance^2^, the metabolically dysregulated state in which increased insulin concentrations are required to maintain euglycemia^3,4^. One leading hypothesis is that as an individual becomes insulin resistant, cells throughout the body–including in the plaque–become dysfunctional and augment atherogenesis. Therefore, it is tempting to speculate that improvements in whole-body insulin sensitivity would reduce atherosclerosis.

On the other hand, a second hypothesis absolves insulin resistance *per se*. Instead, it is directed downstream of insulin resistance, at excessive triglyceride-rich lipoproteins (TRLs). Hypertriglyceridemia is a strong cardiovascular risk factor^5–7^. TRLs are elevated during insulin resistance partly because of hyperinsulinemia and hyperglycemia driving hepatic triglyceride synthesis and secretion, and partly because of impaired TRL clearance^8^. Accordingly, improvements in insulin sensitivity *per se*–without attendant reductions in triglycerides–may be insufficient to reduce atherosclerosis.

To examine whether improving insulin sensitivity is sufficient to reduce atherosclerosis, we leveraged our mouse model lacking hepatic transcription factors FoxO1, FoxO3, and FoxO4. These three proteins are inactivated downstream of insulin signaling, by Akt-mediated phosphorylation and nuclear exclusion^9,10^, and the three isoforms have redundant metabolic functions^11^. A canonical role for hepatic FoxOs is to promote hepatic glucose output^10^. Considerable evidence from multiple investigators, including gold-standard hyperinsulinemic-euglycemic clamps, has demonstrated that ablation of hepatic FoxOs is sufficient to (i) reduce hepatic glucose output, reduce plasma insulin levels, and improve insulin sensitivity^12,13^, and (ii) rescue insulin’s ability to both suppress hepatic glucose production and increase the rate of glucose disposal in mouse models of insulin resistance^14,15^. Thus mice lacking hepatocyte FoxOs are a model of improved systemic insulin sensitivity. On the other hand, ablating FoxOs also boosts triglyceride levels by substantially increasing hepatic de novo lipogenesis^13,16^.

In this work, we used our hepatocyte-specific triple FoxO knockout mice (L-FoxO1,3,4) in two different model systems of atherosclerosis: after injection with gain-of-function Pcsk9 (AAV8.mPcsk9^D377Y^)^17^, and after crossing with Ldlr^−/−^ mice. If improving whole-body insulin sensitivity is sufficient to reduce atherosclerosis, L-FoxO1,3,4 mice should be less susceptible to atherosclerosis. On the other hand, if lowering triglycerides is an imperative mechanism for reducing atherosclerosis, then knocking out FoxOs would be insufficient to lower atherosclerosis.

## Materials and methods

### Mice and diets

All mice were maintained on a 12-hour light/12-hour dark cycle. L-FoxO1,3,4 mice have been described previously^11^. Age-matched littermate controls were homozygous for floxed alleles of FoxO1, FoxO3, and FoxO4, but were Cre-negative. Male mice were injected intravenously with AAV8.mPcsk9^D377Y^ (Addgene 58376, purified at Penn Vector Core)^17^ at a dose of 10^12^ viral genomes per mouse. At the time of viral transduction and diet switch: in cohort 1, the mice were 14.9 ± 0.3 weeks old, and tissues were harvested 12 weeks later; in cohort 2, the mice were 18.6 ± 0.3 weeks old, and tissues were harvested 20 weeks later. L-FoxO1,3,4 mice were also crossed with Ldlr^−/−^ mice (Jackson labs) to generate L-FoxO1,3,4:Ldlr^−/−^, of which both male and female mice were studied. Mice were switched to the western diet, containing 42% kcal from fat and 0.2% cholesterol (Envigo, TD.88137) at 7-8 weeks of age (for Ldlr^−/−^) or at the time of virus injection (for Pcsk9).

### Metabolic tests

Blood glucose was measured by Breeze2 monitor and strips (Bayer). Insulin was measured by ELISA (Millipore Sigma). For glucose tolerance tests, mice were fasted for 16 hours and injected intraperitoneally with 2 g/kg glucose. Cholesterol and triglyceride kits were from Wako. For fast protein liquid chromatography, we used 200 μl of plasma on Superose 6 10/300 GL column (Amersham Pharmacia Biotech) and FC-204 fraction collector (Gilson). For sequential density ultracentrifugation, we separated VLDL (d < 1.006), LDL (1.006 <d <1.063), and HDL (1.063 <d <1.210 g/mL) using NaBr buffers in the Optima MAX-TL Ultracentrifuge with the TLA-100 rotor (Beckman Coulter). Complete blood count analysis was carried out using Heska Element HT5.

### Protein quantitation

Plasma fractions were separated on 4-15% polyacrylamide gradient gels (BioRad). Antibodies were: Ldlr (Abcam Ab30532), β-actin (Cell Signaling Technology, 4970), apoB (Abcam Ab20737), apoA1 (Meridian, K23500R), apoE (Meridian, K23100R), and apoM (LSBio, catalog LS-C319551). Plasma IL-6, IL-10 and TNF-alpha levels were quantified by Milliplex MAP Mouse Cytokine/Chemokine Magnetic Bead Panel (MilliporeSigma, MCYTOMAG-70K).

### Atherosclerotic lesion area

Aortic roots were harvested for histological analysis. Lesional and necrotic analyses were performed on hematoxylin and eosin (H&E)-stained lesional cross-sections and were quantified using an Olympus IX70 with an Olympus DP74 microscope digital camera. Briefly, frozen specimens were immersed in OCT, cryosectioned, and 10µm sections were placed on glass slides. Atherosclerotic lesion area, defined as the region from the internal elastic lamina to the lumen, was quantified by taking the average of 12 sections per mice apart beginning at the base of the aortic root. Quantification was performed in a blinded fashion.

### Statistics

Data are presented as the mean ± SEM. Differences between control and knockout mice were examined using Student’s *t* tests. P<0.05 was considered statistically significant.

## Results

### Improved glucose metabolism and lower insulin in L-FoxO1,3,4:Pcsk9 mice

L-FoxO1,3,4:Pcsk9 mice fed the western diet showed lower fasting glucose and insulin levels than littermate control:Pcsk9 mice (**Figure 1A-B**). L-FoxO1,3,4:Pcsk9 mice also showed substantially better glucose tolerance upon intraperitoneal glucose administration (**Figure 1C**). These effects are consistent with the well-established effect of hepatic FoxOs to promote hepatic glucose output^12,13^, though there may have been a contribution from slightly reduced body weight gain in this cohort (**Supplementary Figure 1A**). Improved glucose metabolism in the setting of lower insulin levels demonstrate that in the atherogenic setting of Pcsk9 gain-of-function expression, hepatic FoxO ablation improves insulin sensitivity. This aligns with considerable evidence from gold-standard hyperinsulinemic-euglycemic clamps^13–15^.

**Figure 1.**
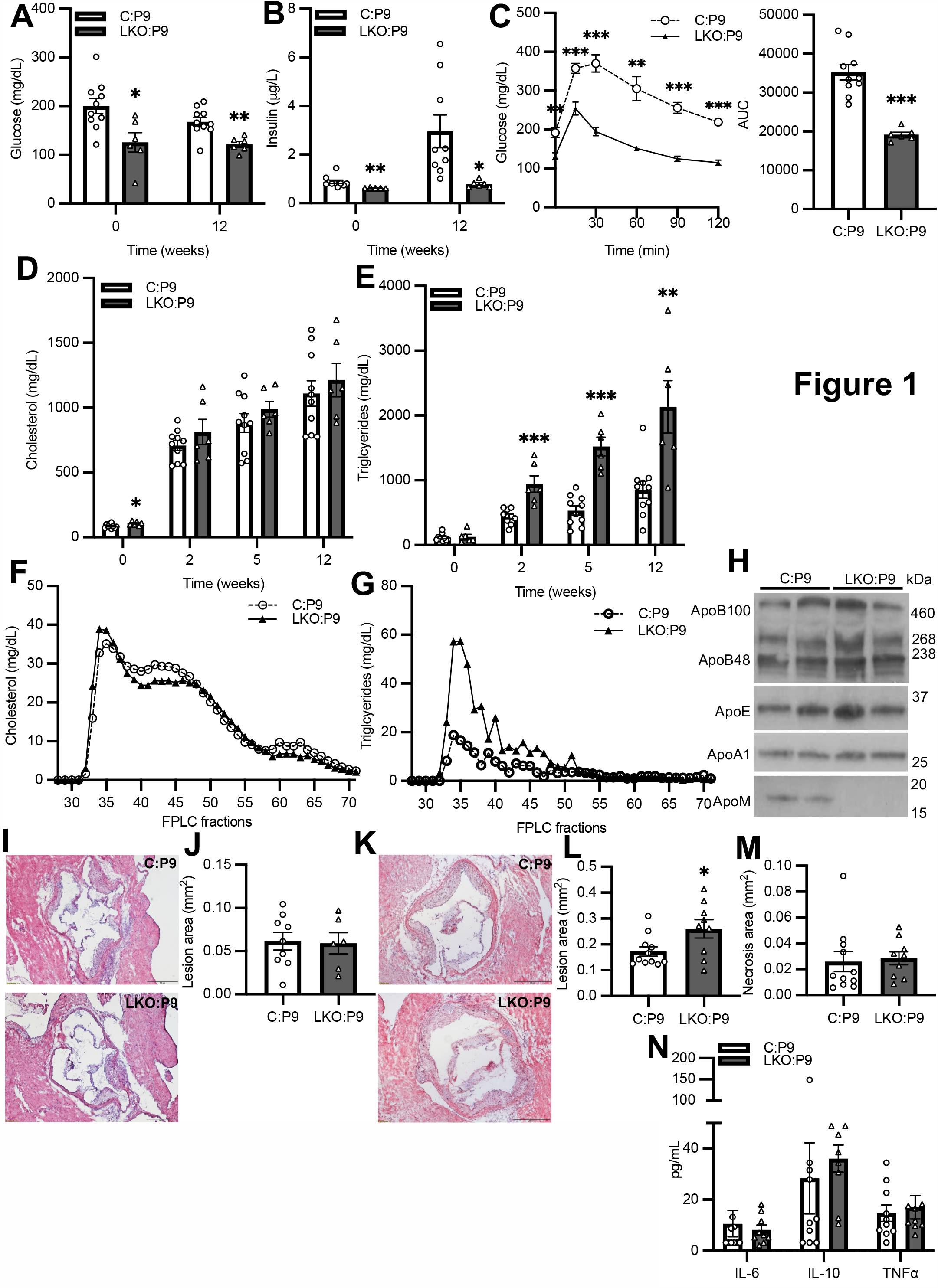
Metabolic parameters and atherosclerotic lesion area in mice injected with Pcsk9 gain-of-function virus. (A-B) Blood glucose and plasma insulin after 5 hours fasting. (C) Intraperitoneal glucose tolerance test. (D-E) Plasma cholesterol and triglyceride levels. (F-G) Cholesterol and triglyceride levels in FPLC-fractionated plasma. (H) Representative western blots of plasma apoB100, apoB48, apoE, apoA1, and apoM. (I) Representative hematoxylin & eosin-stained aortic root sections 12-weeks after western diet feeding. (J) Aortic root lesion area quantified. (K) Representative hematoxylin & eosin-stained aortic root sections 20-weeks after western diet feeding. (L) Aortic root plaque area quantified. (M) Aortic root necrotic core area quantified. (N) Circulating cytokine levels. For A-J, n = 6-10 males/group. For K-N, n = 9-11 males/group. Data are presented as the mean ± SEM. ^*^P < 0.05, ^**^P < 0.01, and ^***^P < 0.001, by Student’s *t* tests. Scale bar represents 500 mm. C:P9, control:Pcsk9 mice; LKO:P9, L-FoxO1,3,4:Pcsk9 mice.

### L-FoxO134:Pcsk9 mice have severe hypertriglyceridemia with WTD feeding

As expected, Pcsk9-injected mice showed depletion of hepatic Ldlr protein (**Supplementary Figure 1B**) and hypercholesterolemia (**Figure 1D**). Pcsk9-injected mice also showed increases in circulating triglycerides, and L-FoxO1,3,4:Pcsk9 mice were strongly disproportionately affected, with 2-3x higher triglycerides than control:Pcsk9 mice (**Figure 1E**). This is consistent with the 2-3x increased rate of hepatic de novo lipogenesis in these mice^13^.

We fractionated plasma by FPLC and found similar cholesterol distributions between genotypes, although in L-FoxO1,3,4:Pcsk9 mice cholesterol was slightly shifted towards larger, VLDL-sized particles (**Figure 1F**). L-FoxO1,3,4:Pcsk9 mice also showed higher triglycerides in all fractions (**Figure 1G**). In whole plasma (**Figure 1H**) or ultracentrifuge-fractionated plasma (**Supplementary Figure 1C**), we observed no major differences in apoB or apoA1. Consistent with our recent finding that FoxOs are necessary for hepatic apoM expression^18^, apoM was nearly absent in whole and fractionated plasma (HDL fraction) from L-FoxO1,3,4:Pcsk9 mice.

### No difference in early atherosclerotic lesion area in L-FoxO1,3,4:Pcsk9 mice

12-weeks after injecting mice with AAV8.mPcsk9^D377Y^ and initiating western diet feeding (*i*.*e*. ∼10 weeks after hypercholesterolemia), we harvested aortic roots and other tissues. We found that L-FoxO1,3,4:Pcsk9 mice displayed higher liver, spleen, and kidney weights (**Supplementary Figure 1D-F**); the causes of these phenotypes are unknown. On the other hand, we found that atherosclerotic lesions were small and not different between genotypes (**Figure 1I-J**).

### Glucose and lipid metabolic phenotypes are reproducible

We carried out a second cohort of Pcsk9-injected mice, to examine a later time point in atherogenesis. We found that key metabolic phenotypes described above were reproducible. L-FoxO1,3,4:Pcsk9 mice showed lower glucose, equivalent hypercholesterolemia, and more severe hypertriglyceridemia than control:Pcsk9 mice (**Supplementary Figure 2A-C**). We observed no differences in apoB levels, though apoA1 was slightly lower and apoM was very low in L-FoxO1,3,4:Pcsk9 mice (**Supplementary Figure 2D**).

**Figure 2.**
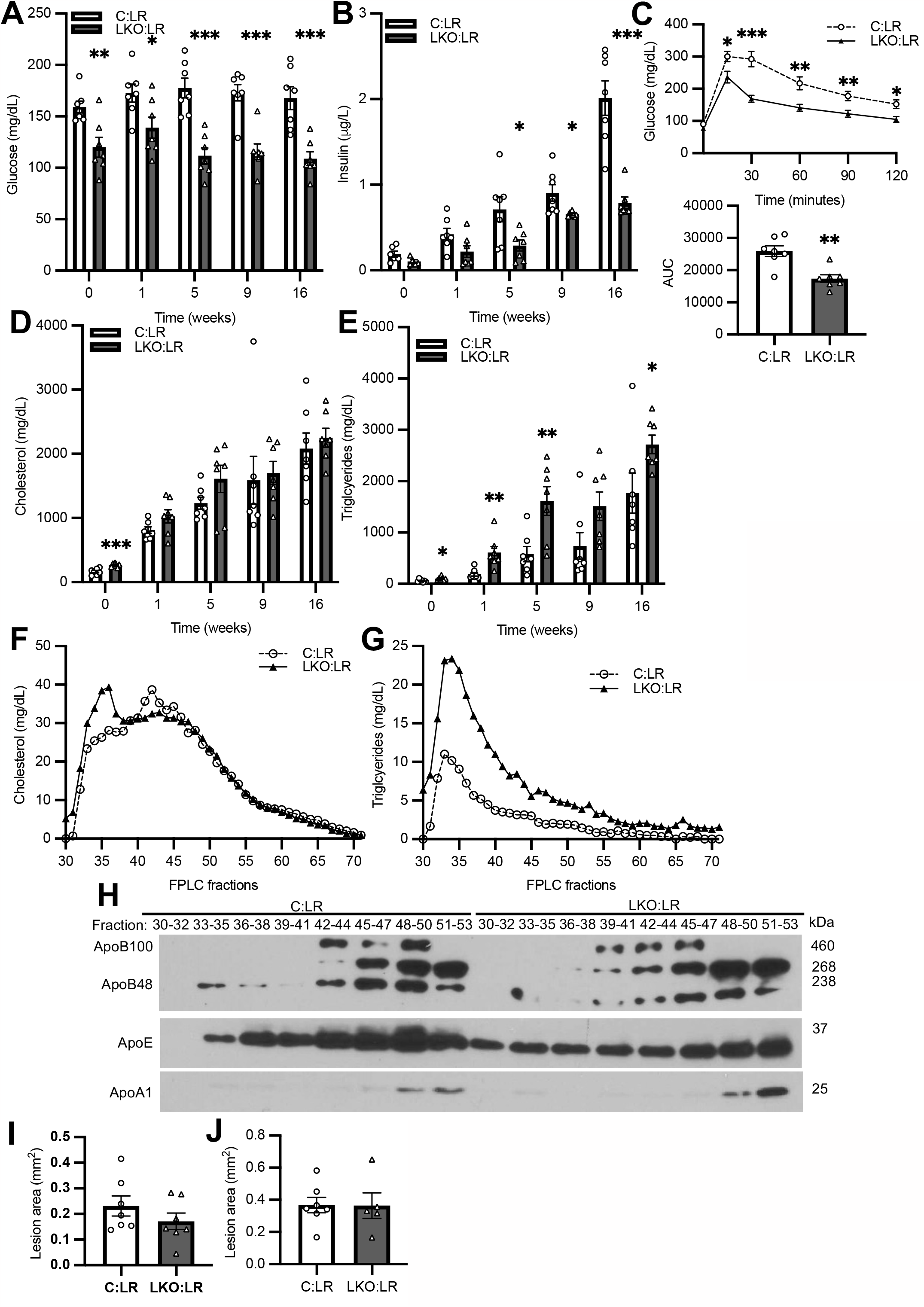
Metabolic parameters and atherosclerosis in Ldlr^−/−^ mice. (A-B) Blood glucose and plasma insulin after 5 hours fasting. (C) Intraperitoneal glucose tolerance test. (D-E) Plasma cholesterol and triglyceride levels. (F-G) Cholesterol and triglyceride levels in FPLC-fractionated plasma. (H) Representative western blots of apoB100, apoB48, apoE, and apoA1 expression in FPLC-fractionated plasma. (I) Aortic root lesion area in males quantified. (J) Aortic root lesion area in females quantified. For A-I, n = 7 males/group. For J, n = 5-7 females/group. Data are presented as the mean ± SEM. ^*^P < 0.05, ^**^P < 0.01, and ^***^P < 0.001, by Student’s *t* tests. C:LR, control:Ldlr^−/−^ mice; LKO:LR, L-FoxO1,3,4:Ldlr^−/−^ mice.

### Increase in late atherosclerotic lesion area in L-FoxO1,3,4:Pcsk9 mice

20-weeks after injecting mice with Pcsk9 and initiating western diet feeding (*i*.*e*. ∼18 weeks after development of hypercholesterolemia), we harvested tissues. L-FoxO1,3,4:Pcsk9 mice displayed higher liver, spleen, and kidney weights as we observed previously in the earlier cohort (**Supplementary Figure 2E-G**). At this 20-week time point, atherosclerotic lesion area was 50% larger in L-FoxO1,3,4:Pcsk9 mice compared to control:Pcsk9 mice (**Figure 1K-L**). There were no significant differences in necrotic core size or circulating cytokines IL-6, IL-10, or Tnfα (**Figure 1M-N**).

### Improved insulin sensitivity in male L-FoxO1,3,4:Ldlr^−/−^ mice

We used Ldlr-deficiency as a second model of atherosclerosis. Male L-FoxO1,3,4:Ldlr^−/−^ mice showed lower fasting glucose and insulin compared to littermate control:Ldlr^−/−^ mice (**Figure 2A-B**), despite similar body weight and body composition (**Supplementary Figure 3A-C**). L-FoxO1,3,4:Ldlr^−/−^ mice showed markedly improved intraperitoneal glucose tolerance (**Figure 2C**). The combination of improved glucose metabolism with lower insulin levels again demonstrates improved insulin sensitivity in mice lacking hepatic FoxOs in the atherogenic setting of Ldlr-deficiency.

### L-FoxO134:Ldlr^-/-^ mice have severe hypertriglyceridemia with WTD feeding

After switching to western diet, all Ldlr-deficient mice developed equivalent hypercholesterolemia (**Figure 2D**). On the other hand, L-FoxO1,3,4:Ldlr^−/−^ mice showed higher triglycerides throughout the experiment (**Figure 2E**). By FPLC fractionation, we observed that in L-FoxO1,3,4:Ldlr^−/−^ mice, cholesterol was shifted to larger, VLDL-sized particles and triglycerides were higher in all fractions (**Figure 2F-G**). This was accompanied by a slight leftward shift of apoB elution (**Figure 2H**).

### No difference in atherosclerotic lesion area in male L-FoxO1,3,4:Ldlr^−/−^ mice

16-weeks after western diet initiation, we harvested tissues. Male L-FoxO1,3,4:Ldlr^−/−^ mice showed higher liver, spleen, and kidney weights compared to control Ldlr^−/−^ mice (**Supplementary Figure 3D-F**). There were no differences between genotypes in complete blood count analysis (**Supplementary Figure 3G**). We observed no significance differences between genotypes in atherosclerotic lesion area (**Figure 2I**).

### Mild metabolic phenotypes and no difference in atherosclerotic lesion area in female L-FoxO1,3,4:Ldlr^−/−^ mice

Female L-FoxO1,3,4:Ldlr^−/−^ mice showed lower or normal fasting glucose and insulin, compared to female littermate Ldlr^−/−^ controls, and both genotypes gained similar amounts of body weight (**Supplementary Figure 4A-C**). Mice of both genotypes were equally hypercholesterolemic, and although hypertriglyceridemia developed faster in female L-FoxO1,3,4:Ldlr^−/−^ mice, both genotypes ultimately developed equivalent triglycerides (**Supplementary Figure 4D-E**). 16-weeks after western diet initiation, we harvested tissues. Female L-FoxO1,3,4:Ldlr^−/−^ mice showed higher liver weights compared to control Ldlr^−/−^ mice (**Supplementary Figure 4F**). However, we observed no significance differences in atherosclerotic lesion area between genotypes (**Figure 2J**).

## Discussion

Type 2 diabetes is associated with increased atherosclerosis. One contributor is hyperglycemia, though this does not fully explain the increased cardiovascular risk in individuals with diabetes^19^. Another leading candidate to explain the increased atherosclerosis is insulin resistance^20^. Insulin signaling occurs in cells and tissues throughout the body, and cellular insulin resistance has been reported to negatively affect the functions of multiple atherosclerosis-relevant cell types such as macrophages^21^, endothelial cells^22^, and vascular smooth muscle cells^23^. This raises the possibility that improving insulin sensitivity would be atheroprotective.

In this work, we demonstrate that improving insulin sensitivity in mice by knocking out hepatic FoxOs is insufficient to lower atherosclerosis. In at least one setting (20 weeks after Pcsk9 injection), hepatic FoxO knockout mice actually showed larger atherosclerotic lesions. Because loss of hepatic FoxOs potently improves glucose metabolism and whole-body insulin sensitivity, our findings suggest that systemic insulin sensitization is not anti-atherogenic.

There are other compelling explanations for the link between insulin resistance and atherosclerosis. One possibility is that a third factor, such as inflammation^27^ or oxidative stress^28^ underlies both insulin resistance and atherosclerosis. But a second possibility calls upon TRLs^24^. TRLs are strong cardiovascular risk factors^5,6^, and they are elevated in individuals with insulin resistance and type 2 diabetes^25^. Because hepatic FoxOs have the dual effect of promoting glucose production and suppressing lipogenesis, loss of hepatic FoxOs results in decreased glucose production, but increased lipogenesis and increased VLDL secretion^13,16,26^. Thus in our mouse model, insulin sensitization is necessarily coincident with increased triglycerides. A potential interpretation of our findings is that insulin sensitization may only be anti-atherogenic if TRLs are effectively lowered.

Overall, this study does not support a direct effect of insulin sensitivity on atherogenesis. However, it is worth noting that we investigated early and moderate time points of atherosclerotic lesion development. We cannot exclude the possibility that insulin sensitization may have beneficial effects on advanced lesions^29^ or on atherosclerosis regression^8^, and it will be of interest to investigate those in the future.

## Acknowledgements

MCI, NS, KZ, and MH performed experiments and analyzed data. LO, GF and RH supervised experiments and analyzed data. MH and RH wrote the manuscript. RH had full access to all the data in the study and takes responsibility for its integrity and the data analysis. All authors edited the manuscript. We are grateful to Thomas Kolar for technical assistance and to our colleagues at Columbia and Albany Medical College for helpful discussions.

## Supplemental material

Supplementary Figure 1

Supplementary Figure 2

Supplementary Figure 3

Supplementary Figure 4

## REFERENCES

1. Tsao CW, Aday AW, Almarzooq ZI, Alonso A, Beaton AZ, Bittencourt MS, Boehme AK, Buxton AE, Carson AP, Commodore-Mensah Y, Elkind MSV, Evenson KR, Eze-Nliam C, Ferguson JF, Generoso G, Ho JE, Kalani R, Khan SS, Kissela BM, Knutson KL, Levine DA, Lewis TT, Liu J, Loop MS, Ma J, Mussolino ME, Navaneethan SD, Perak AM, Poudel R, Rezk-Hanna M, Roth GA, Schroeder EB, Shah SH, Thacker EL, VanWagner LB, Virani SS, Voecks JH, Wang NY, Yaffe K, Martin SS, null null. Heart Disease and Stroke Statistics—2022 Update: A Report From the American Heart Association. Circulation. 2022;145(8):e153–e639. doi:10.1161/CIR.0000000000001052

2. Reaven G. Insulin Resistance and Coronary Heart Disease in Nondiabetic Individuals. Arteriosclerosis, Thrombosis, and Vascular Biology. 2012;32(8):1754–1759. doi:10.1161/ATVBAHA.111.241885

3. James DE, Stöckli J, Birnbaum MJ. The aetiology and molecular landscape of insulin resistance. Nat Rev Mol Cell Biol. 2021;22(11):751–771. doi:10.1038/s41580-021-00390-6

4. Reaven GM. Banting lecture 1988. Role of insulin resistance in human disease. Diabetes. 1988;37:1595–1607.

5. Bansal S, Buring JE, Rifai N, Mora S, Sacks FM, Ridker PM. Fasting compared with nonfasting triglycerides and risk of cardiovascular events in women. JAMA. 2007;298:309–316. doi:298/3/309 [pii] 10.1001/jama.298.3.309 [doi]

6. Nordestgaard BG, Benn M, Schnohr P, Tybjaerg-Hansen A. Nonfasting triglycerides and risk of myocardial infarction, ischemic heart disease, and death in men and women. JAMA. 2007;298:299–308. doi:298/3/299 [pii] 10.1001/jama.298.3.299 [doi]

7. Sandesara PB, Virani SS, Fazio S, Shapiro MD. The Forgotten Lipids: Triglycerides, Remnant Cholesterol, and Atherosclerotic Cardiovascular Disease Risk. Endocr Rev. 2019;40(2):537–557. doi:10.1210/er.2018-00184

8. Eckel RH, Bornfeldt KE, Goldberg IJ. Cardiovascular disease in diabetes, beyond glucose. Cell Metabolism. 2021;33(8):1519–1545. doi:10.1016/j.cmet.2021.07.001

9. Haeusler RA, McGraw TE, Accili D. Biochemical and cellular properties of insulin receptor signalling. Nat Rev Mol Cell Biol. Published online October 4, 2017. doi:10.1038/nrm.2017.89

10. Lin HV, Accili D. Hormonal regulation of hepatic glucose production in health and disease. Cell Metab. 2011;14:9–19. doi:S1550-4131(11)00221-X [pii] 10.1016/j.cmet.2011.06.003 [doi]

11. Haeusler RA, Kaestner KH, Accili D. FoxOs Function Synergistically to Promote Glucose Production. J Biol Chem. 2010;285:35245–35248. doi:C110.175851 [pii] 10.1074/jbc.C110.175851 [doi]

12. Matsumoto M, Pocai A, Rossetti L, Depinho RA, Accili D. Impaired regulation of hepatic glucose production in mice lacking the forkhead transcription factor Foxo1 in liver. Cell Metab. 2007;6:208–216.

13. Haeusler RA, Hartil K, Vaitheesvaran B, Arrieta-Cruz I, Knight CM, Cook JR, Kammoun HL, Febbraio MA, Gutierrez-Juarez R, Kurland IJ, Accili D. Integrated control of hepatic lipogenesis versus glucose production requires FoxO transcription factors. Nature communications. 2014;5:5190. doi:10.1038/ncomms6190

14. O-Sullivan I, Zhang W, Wasserman DH, Liew CW, Liu J, Paik J, DePinho RA, Stolz DB, Kahn CR, Schwartz MW, Unterman TG. FoxO1 integrates direct and indirect effects of insulin on hepatic glucose production and glucose utilization. Nat Commun. 2015;6:7079. doi:10.1038/ncomms8079

15. Lu M, Wan M, Leavens KF, Chu Q, Monks BR, Fernandez S, Ahima RS, Ueki K, Kahn CR, Birnbaum MJ. Insulin regulates liver metabolism in vivo in the absence of hepatic Akt and Foxo1. Nat Med. 2012;18:388–395. doi:nm.2686 [pii] 10.1038/nm.2686 [doi]

16. Tao R, Wei D, Gao H, Liu Y, DePinho RA, Dong XC. Hepatic FoxOs regulate lipid metabolism via modulation of expression of the nicotinamide phosphoribosyltransferase gene. J Biol Chem. 2011;286:14681–14690. doi:M110.201061 [pii] 10.1074/jbc.M110.201061 [doi]

17. Bjørklund Martin Mæng, Hollensen Anne Kruse, Hagensen Mette Kallestrup, Dagnæs-Hansen Frederik, Christoffersen Christina, Mikkelsen Jacob Giehm, Bentzon Jacob Fog. Induction of Atherosclerosis in Mice and Hamsters Without Germline Genetic Engineering. Circulation Research. 2014;114(11):1684–1689. doi:10.1161/CIRCRESAHA.114.302937

18. Izquierdo MC, Shanmugarajah N, Lee SX, Kraakman MJ, Westerterp M, Kitamoto T, Harris M, Cook JR, Gusarova GA, Zhong K, Marbuary E, O-Sullivan I, Rasmus NF, Camastra S, Unterman TG, Ferrannini E, Hurwitz BE, Haeusler RA. Hepatic FoxOs link insulin signaling with plasma lipoprotein metabolism through an apolipoprotein M/sphingosine-1-phosphate pathway. J Clin Invest. Published online February 1, 2022:e146219. doi:10.1172/JCI146219

19. Ozkan B, Ndumele CE. Addressing Cardiovascular Risk in Diabetes: It’s More Than the Sugar. Circulation. 2023;147(25):1887–1890. doi:10.1161/CIRCULATIONAHA.123.065090

20. Bornfeldt K, Tabas I. Insulin Resistance, Hyperglycemia, and Atherosclerosis. Cell Metabolism. 2011;14:575–585.

21. Tabas I, Tall A, Accili D. The impact of macrophage insulin resistance on advanced atherosclerotic plaque progression. Circ Res. 2010;106:58–67. doi:106/1/58 [pii] 10.1161/CIRCRESAHA.109.208488 [doi]

22. Rask-Madsen C, Li Q, Freund B, Feather D, Abramov R, Wu IH, Chen K, Yamamoto-Hiraoka J, Goldenbogen J, Sotiropoulos KB, Clermont A, Geraldes P, Dall’Osso C, Wagers AJ, Huang PL, Rekhter M, Scalia R, Kahn CR, King GL. Loss of insulin signaling in vascular endothelial cells accelerates atherosclerosis in apolipoprotein E null mice. Cell Metabolism. 2010;11:379–389. doi:10.1016/j.cmet.2010.03.013

23. Li Q, Fu J, Xia Y, Qi W, Ishikado A, Park K, Yokomizo H, Huang Q, Cai W, Rask-Madsen C, Kahn CR, King GL. Homozygous receptors for insulin and not IGF-1 accelerate intimal hyperplasia in insulin resistance and diabetes. Nat Commun. 2019;10(1):4427. doi:10.1038/s41467-019-12368-2

24. Tall AR, Thomas DG, Gonzalez-Cabodevilla AG, Goldberg IJ. Addressing dyslipidemic risk beyond LDL-cholesterol. J Clin Invest. 2022;132(1). doi:10.1172/JCI148559

25. Krauss RM. Lipids and lipoproteins in patients with type 2 diabetes. Diabetes Care. 2004;27:1496–1504.

26. Haeusler RA, Han S, Accili D. Hepatic FoxO1 Ablation Exacerbates Lipid Abnormalities during Hyperglycemia. J Biol Chem. 2010;285:26861–26868. doi:M110.134023 [pii] 10.1074/jbc.M110.134023 [doi]

27. Wu H, Ballantyne CM. Metabolic Inflammation and Insulin Resistance in Obesity. Circulation Research. Published online 2020. doi:10.1161/CIRCRESAHA.119.315896

28. Huang J, Tao H, Yancey PG, Leuthner Z, May-Zhang LS, Jung JY, Zhang Y, Ding L, Amarnath V, Liu D, Collins S, Davies SS, Linton MF. Scavenging dicarbonyls with 5’-O-pentyl-pyridoxamine increases HDL net cholesterol efflux capacity and attenuates atherosclerosis and insulin resistance. Mol Metab. 2023;67:101651. doi:10.1016/j.molmet.2022.101651

29. Han S, Liang CP, DeVries-Seimon T, Ranalletta M, Welch CL, Collins-Fletcher K, Accili D, Tabas I, Tall AR. Macrophage insulin receptor deficiency increases ER stress-induced apoptosis and necrotic core formation in advanced atherosclerotic lesions. Cell Metab. 2006;3:257–266.

